# Microglial cGAS deletion protects against amyloid-β induced Alzheimer’s disease pathogenesis

**DOI:** 10.1101/2023.08.07.552300

**Authors:** Sijia He, Xin Li, Namrata Mittra, Anindita Bhattacharjee, Hu Wang, Shangang Zhao, Feng Liu, Xianlin Han

## Abstract

Innate immune activation plays a vital role in the development of Alzheimer’s disease (AD) and related dementias (ADRD). Among which, the DNA sensing cyclic GMP-AMP synthase (cGAS)- STING pathway has been implicated in diverse aspects of AD progression.

In the current study, we showed that the cGAS-STING signaling was up-regulated in AD and this elevation was mainly contributed by the microglial population other than non-microglial cell types in the brain. By establishing an inducible, microglia-specific cGAS knockout mouse model in 5xFAD background, we found that deleting microglial cGAS at the onset of amyloid-β (Aβ) pathology significantly limited plaque formation, and protected mice from Aβ-induced cognitive impairment. Mechanistically, we found cGAS was necessary for plaque-associated microglial enrichment potentially driven by IRF8, and was indispensable for the development of disease-associated microglia (DAM) phenotype. Meanwhile, the loss of microglial cGAS reduced the levels of dystrophic neurites which led to preserved synaptic integrity and neuronal function. Our study provides new insights in understanding the effects of innate immune in AD via a cell-type specific manner, and lays the foundation for potential targeted intervention of the microglial cGAS-STING pathway toward the improvement of AD.

## 1. Introduction

Alzheimer’s disease (AD), a devastating condition that impairs memory and cognitive function, is becoming increasingly prevalent among the elderly (1). Despite significant research efforts to understand the underlying causes of AD/ADRD (Alzheimer’s disease and related dementias), no definitive cure has been discovered yet. Genome-wide associated studies (GWASs) have identified a plethora of AD risk factors, which include triggering receptor-expressed on myeloid cells 2 (TREM2), apolipoprotein E (APOE), ATP-binding cassette transporter (ABCA) family, CD33, and complement pathway genes (2-4). Intriguingly, more than half of the risk loci that clearly involve a specific gene are significantly enriched or uniquely expressed in immune cells, particularly microglia and macrophages (5). This intriguing finding suggests that innate immune molecules play a pivotal role in the development and progression of the disease (5).

The cGAS (encoded by *Mb21d1* gene)-Stimulator of Interferon Genes (STING) pathway is an essential innate immune signaling for the initiation of type I interferon response to defend against pathogen infections via sensing of double-stranded DNA (dsDNA). Activated by the binding of self- or non-self dsDNAs, cGAS catalyzes the production of 2’3’ cyclic GMA-AMP (2’3’-cGAMP), which in turn acts as a second messenger to activate STING. The activation of STING further promotes interferon gene expression by inducing the TBK1-IRF3-NFκB axis (6, 7). In addition to its classical function in innate immunity against pathogen infection, the involvement of cGAS-STING pathway has been reported in various physiological and pathological processes such as autoimmune (8), cellular senescence (9), autophagy (10), inflammation (11), and cancer immunity (12). Multiple studies have also emphasized its roles in neurological disorders, including multiple sclerosis (13), Parkinson’s disease (14), amyotrophic lateral sclerosis (15), traumatic brain injury (16), stroke (17), and aging-related neurodegeneration (18). Particularly, recent studies suggested a contribution of cGAS-STING pathway activation in the development of AD (19, 20), underscoring the significance of better understanding the roles of cGAS in AD pathogenesis, as well as the potential therapeutic value by targeting the cGAS-STING pathway in treatment of AD.

Despite highly enriched in immune cells such as macrophages and microglia in the brain, multiple lines of evidence have demonstrated the diverse roles of cGAS in non-immune cells. cGAS expression in central nervous system (CNS) neurons and peripheral nervous system (PNS) Schwann cells promotes spontaneous axon regeneration (21). Striatal neuronal cGAS-STING signaling promotes inflammatory and autophagic responses in Huntington disease (22). Expression of cGAS was found in human astrocytes to enable a cellular response to foreign dsDNA (23). Endothelial activation of the cGAS-STING pathway plays a vital role in aging-related endothelial dysfunction (24). These findings suggest a wide distribution of cGAS expression and the potential existence of cell type-specific roles mediated by the cGAS-STING pathway. Further, these cell type-specific effects may act collectively in a diverse or synergistic manner in disease progression. Thus, identifying cell-type specific function is critical for understanding AD pathogenesis and future development of targeted therapeutic strategy.

In the current study, we established an inducible, microglia-specific cGAS knockout model, and investigated its effects in the 5xFAD mouse model of Aβ pathology (which harbors transgenes of amyloid precursor protein (APP) and PSEN1 with five AD-linked mutations). We identified microglia as the major cell type responsible for the up-regulation of cGAS-STING signaling under Aβ pathology. Deletion of microglial cGAS in adult mouse brain ameliorated Aβ plaque formation, and protected mice from Aβ accumulation-induced cognitive impairment. Further, we found cGAS is important for the development of DAM and dystrophic neurites under Aβ pathology. Our findings provide valuable information in understanding the novel role of microglia-specific cGAS in modulating pathogenesis of AD.

## 2. Results

### 2.1 Microglia are the main contributors to the up-regulated cGAS-STING pathway in the AD brain

In our investigation of cGAS in AD, we initially assessed its expression in postmortem AD patient tissues and normal controls. As previously reported (19), cGAS protein levels were found to be elevated in the AD brain compared to controls (Fig. 1a). Consistent with this finding, we observed an increase in cGAS mRNA levels in AD spinal cord tissues (Fig. 1b). While cGAS expression has been traditionally associated with immune cells (7), recent studies have suggested its involvement in non-immune cells in the brain, such as neurons (21) and astrocytes (23). Consistently, our analysis of human brain tissue data from online resources (25, 26) demonstrated high cGAS expression in microglia, moderate expression in neurons and oligodendrocytes, and detectable but lower levels in astrocytes and endothelial cells (Fig. S1a-c). Similar expression patterns were observed in spinal cord tissue based on the data from a single cell sequencing study (27) (Fig. S1d).

**Figure 1.**
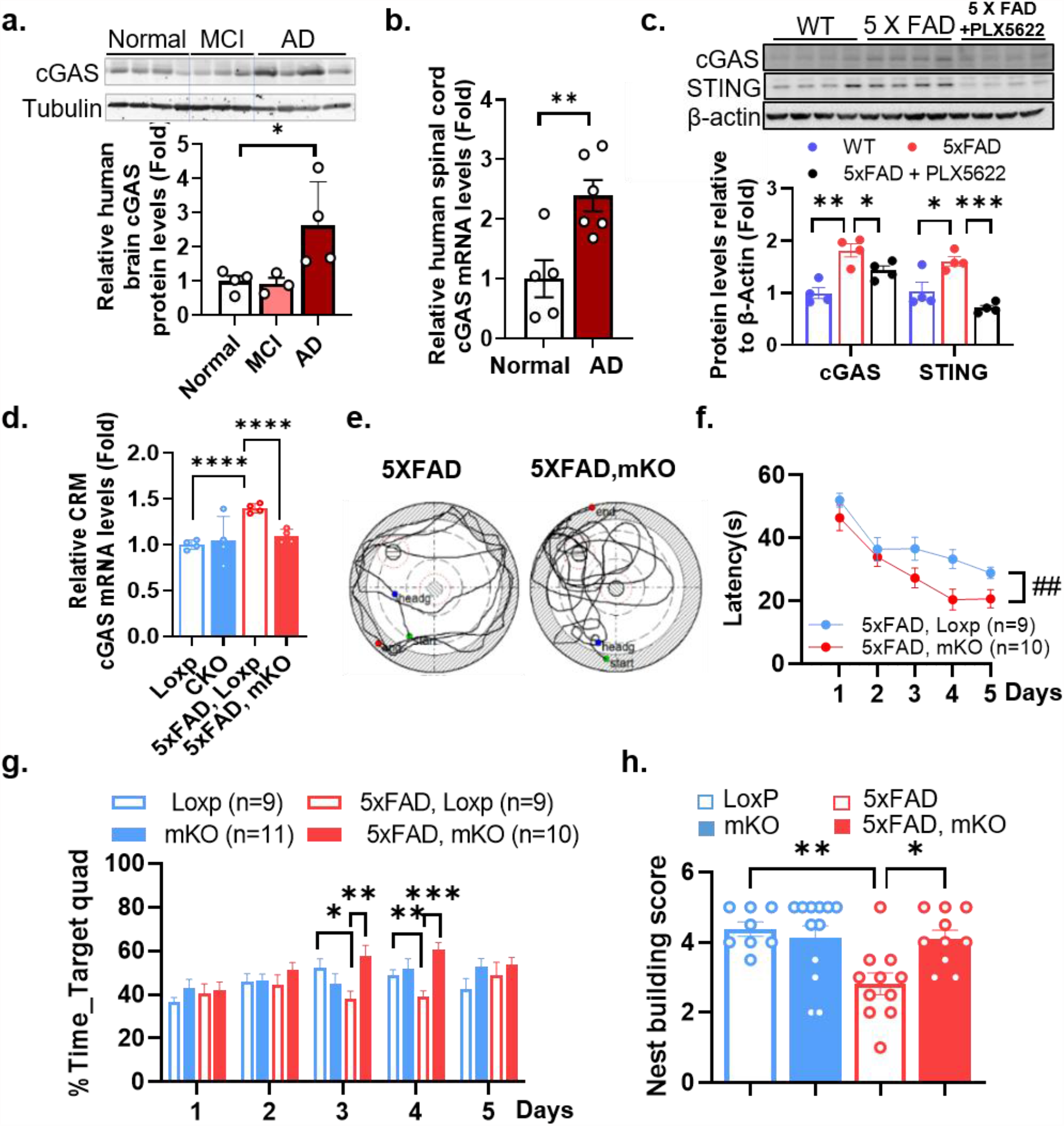
Microglia-specific cGAS deletion protects mice from Aβ accumulation-induced cognitive deficits. a) Western-blot evaluation of cGAS protein levels in human postmortem brain samples. MCI: mild cognitive impairment. n=3-4/group. b) qPCR measurement of cGAS mRNA levels from human spinal cord samples of normal and AD individuals. n=5-7/group. c) Protein levels of cGAS and STING in the mouse brain lysate. (Female, 9 months age, n=4/group). d) Cerebrum tissue mRNA levels of cGAS measured with qPCR. n=4/genotype. e) Representative tracking, f) Latency, and g) Percent time in target quadrant during training in MWM test. (Showing male data from 6 months old). h) Nest building test score. (Male and female, 7 months old, n=8-12/group). Student’s t - test for b), ANOVA followed with Tukey’s test for a), c), d), g), and f). *p ≤ 0.05, **p ≤ 0.01, ***p ≤ 0.001, and ****p ≤ 0.0001.

To determine the cell population responsible for cGAS elevation in AD, we treated 5xFAD mice with the CSF1R inhibitor PLX5622, known to selectively deplete microglia (28). In line with the induction of cGAS signaling observed in human AD, both cGAS and STING protein levels were significantly induced in 5xFAD mice compared to age-matched wild-type controls (Fig. 1c). Interestingly, the induction was abolished upon microglial elimination (Fig. 1c), suggesting that microglia are the primary contributors to the elevation of cGAS in AD.

### 2.2 Microglia-specific cGAS deletion protects mice from Aβ accumulation-induced cognitive deficits

To explore the potential role of cGAS up-regulation in the process of AD, we generated a tamoxifen (TM)-inducible, microglia-specific cGAS knockout mouse line (mKO) by crossing Cx3cr1-Cre^ERT2^ with cGAS flox/flox mice (Loxp) (Fig. S2a). The mKO was further crossed with 5xFAD mice to generate control and conditional knockout mice in an Aβ pathology background (i.e., 5xFAD, and 5xFAD-mKO). To avoid any developmental disruption, we injected mice with TM (80 mg/kg bodyweight for 4 consecutive days) at 2 months of age to induce *Mb21d1* gene deletion in adult microglial populations. qPCR detected the induction of cGAS mRNA in 5xFAD vs. Loxp, while this induction was abolished in 5xFAD-mKO mouse cerebrum (CRM) (Fig. 1d), reflecting a successful establishment of the cGAS mKO model. To characterize the effect of microglial cGAS deletion on cognitive function, we performed behavioral tests at 4 months post TM induction. Of special interest, mice lacking cGAS (5xFAD-mKO) reached the hidden platform with significantly lower latency than 5xFAD littermates in a Morris water maze (MWM) test (Fig. 1e, 1f, and Fig. S2b). This was also reflected by the increased time they spent in the target quadrant (Fig. 1g and Fig. S2c). Furthermore, deletion of microglial cGAS greatly improved performance in a nest building test (Fig. 1h and Fig. S2d). Together, these results established a protective effect in the absence of microglial cGAS, suggesting that the induction of cGAS in Aβ pathology is a strong driving factor for cognitive decline in AD.

### 2.3 Selective microglial cGAS ablation significantly decreases the plaque load in 5xFAD mouse brains

In light of the improved behavior in 5xFAD-mKO, we wondered if it was associated with altered Aβ pathology. By staining plaque with an anti-Aβ peptide antibody, we found a considerable decrease of amyloid deposits overall in both male and female 5xFAD-mKO vs. 5xFAD littermates (Fig. 2a and 2b). A marked decrease of fibrillary Aβ in 5xFAD-mKO brains was also observed by methoxy-x04 staining (Fig. 2c and 2d). Consistent with these findings, Western-blot evaluation showed significant decline of Aβ peptide levels in the CRM lysate of 5xFAD-mKO vs. 5xFAD (Fig. 2e). Additionally, both Aβ42 and Aβ40 levels were markedly decreased in soluble and insoluble fractions (Fig. 2f and 2g), while Aβ38 was decreased in soluble fraction (Fig. 2h), as determined by enzyme-linked immunosorbent assay (ELISA). These observations suggest that microglial cGAS may play an important role in facilitating Aβ accumulation in AD pathology.

**Figure 2.**
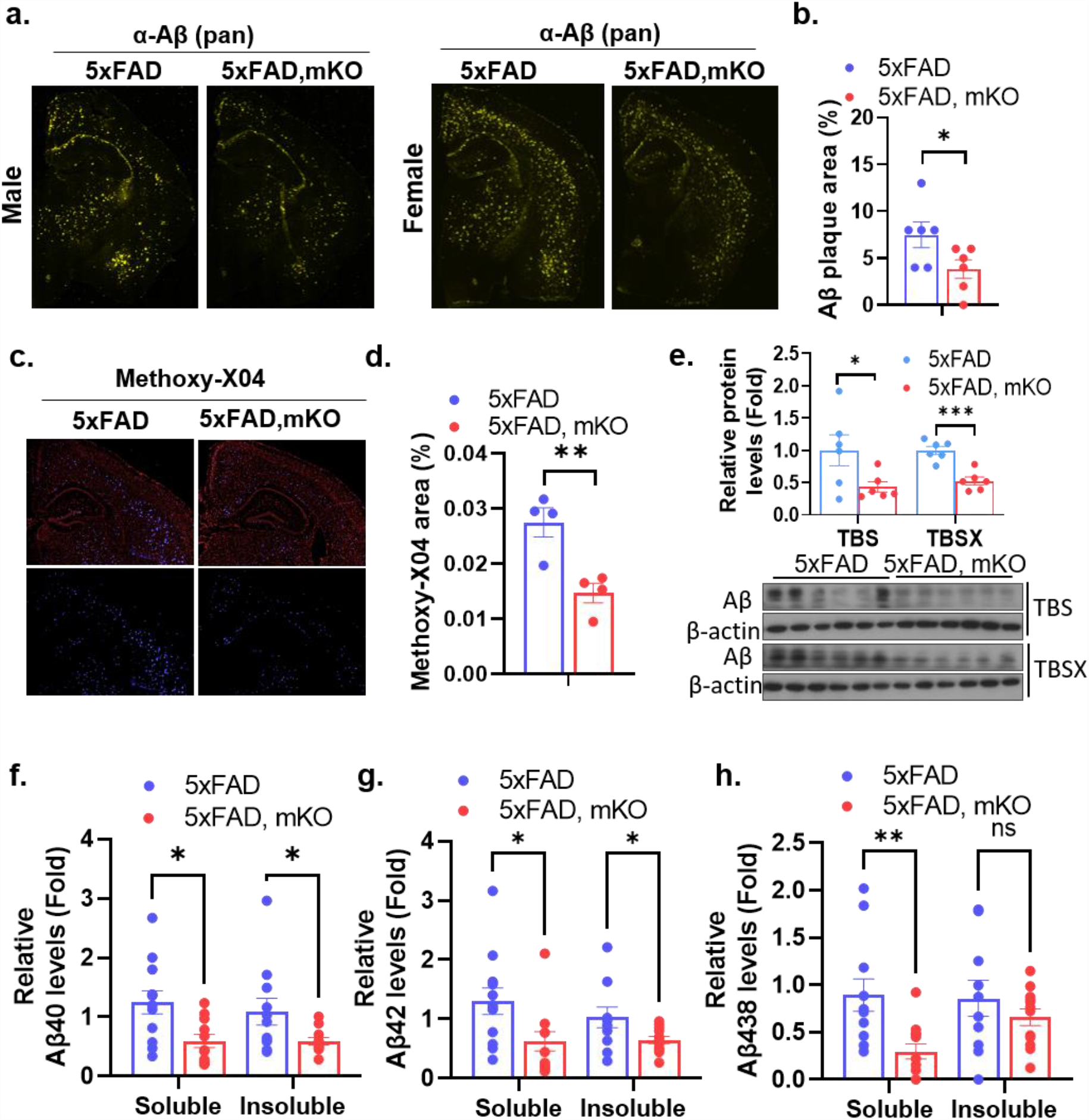
Selective microglial cGAS ablation significantly decreases plaque loads in 5xFAD mouse brains. a) Immunofluorescence (IF) staining of mouse cerebrum sections with a pan Aβ antibody to evaluate Aβ accumulation. b) Quantification of a), n=6/group, male and female combined. c) Evaluation of plaque loads using Methoxy-X04 staining (Male). d) Quantification of c), n=4/group. e) Western-blot evaluation of Aβ levels in male cerebral TBS and TBSX lysates, n=6/group. f) Aβ40, g) Aβ42, and h) Aβ38 levels in mouse brain lysates measured with ELISA. n=11-12/group, male and female combined. Soluble fraction represents protein levels in TBS and TBSX combined equally, insoluble fraction represents protein levels from GuHCl isolation. Student’s t-test for b) and d). ANOVA followed with Tukey’s test for e) - h). *p ⩽ 0.05, **p ⩽ 0.01, and ns (p > 0.05).

### 2.4 Conditional cGAS deletion blunts microglial responses to Aβ pathology

We next asked whether cGAS reduction affects microglial activation in response to Aβ. Co-staining with Iba1, DAPI and Aβ antibodies revealed similar numbers of DAPI^+^ microglia in the hippocampal area of 5xFAD-mKO and 5xFAD mice, while the numbers of microglia within a 10-μm radius from plaque borders were significantly reduced in 5xFAD-mKO compared to 5xFAD (Fig. 3a-c), suggesting decreased microglial reactivity in response to plaque deposition. Similar observation was also found in the cortex region of the brain (Fig. 3d-f). The decline of plaque associated microglia was further evidenced by a decreased number of PU.1^+^ nuclei detected in a 10-μm radius from methoxy-x04^+^ plaque borders in 5xFAD-mKO vs. 5xFAD littermates (Fig. 3g and 3h). Of note, no differences in microglial numbers or Iba1 staining were found when comparing mKO and Loxp mice without amyloid pathology (Fig. 3i), suggesting that cGAS appears to be redundant for maintaining microglial numbers and activity in the steady state. These observations indicated that the presence of cGAS signaling is necessary for Aβ plaque-induced microglial responses.

**Figure 3.**
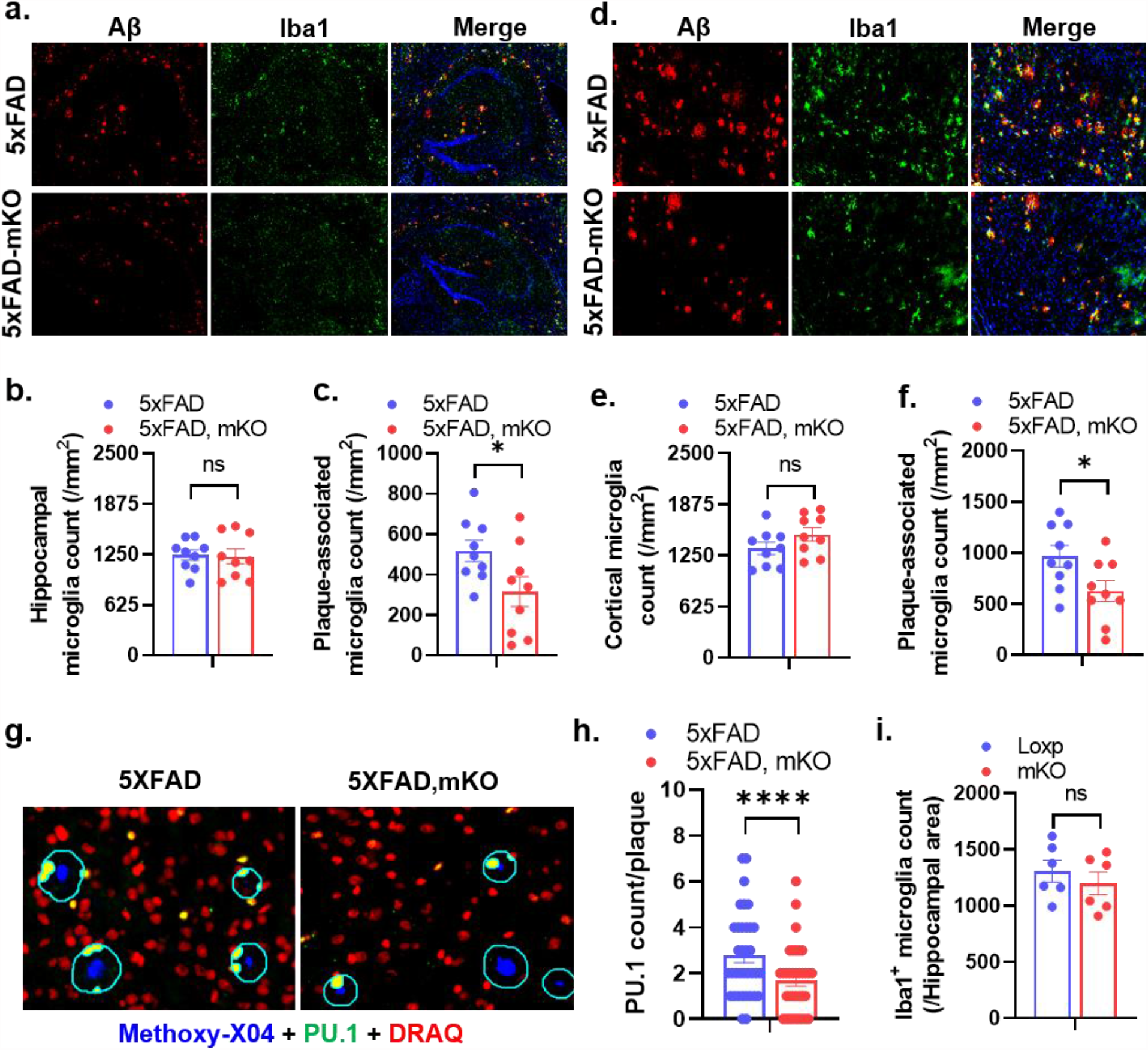
cGAS ablation ameliorates microgliosis in 5xFAD mouse brains. a) Hippocampal IF co-staining using Aβ and Iba1 antibodies with DAPI (blue color) in 7-months-old mice. b) Quantification of overall hippocampal microglial density, and c) Plaque-associated microglial density in the hippocampus area, n=3 mice/group with 3 sections for each mouse. d) Cortical IF co-staining using Aβ and Iba1 antibodies with DAPI. e) Quantification of overall cortical microglial density, and f) Plaque-associated microglia density in cortex area, n=3 mice/group with 3 sections for each mouse. g) Analysis of plaque-associated microglia count within 10 nm distance to plaques using Methoxy-X04 co-staining with PU.1 and DRAQ (DNA stain indicating nuclei). h) Quantification of g), n=35 plaques/genotype. i) Quantification of Iba1 IF staining in Loxp and mKO mice without plaques. Student’s t-test *p ⩽ 0.05, ****p ⩽ 0.0001, and ns (p > 0.05).

### 2.5 Loss of microglial cGAS attenuates microgliosis and ameliorates the generation of disease-associated microglia

Neuroinflammation is one of the cardinal features of AD. Activation of microglia in response to Aβ elicits expression of inflammatory and anti-inflammatory cytokines, which play multifarious roles in neurodegeneration and neuroprotection (29). Given the reduced microglial response to plaques in the microglial cGAS-deficient mice, we attempted to understand the mechanistic regulation of such a phenotype. We evaluated gene expression profile of mouse brain tissue using the NanoString neuroinflammation panel. Principle component analysis (PCA) showed a clear separation of male 5xFAD group compared to groups without Aβ pathology (Loxp and mKO), while the loss of cGAS in a 5xFAD background led to an overlap of 95% confidence interval with Loxp controls (Fig. 4a). An overall pathway analysis indicated a drastically altered functional pathways between 5xFAD-mKO and 5xFAD, most significantly-impacted pathways were related to immune and inflammation regulation (Fig. 4b). We also detected marked changes in pathways including astrocyte function, autophagy, and lipid and carbohydrate metabolism (Fig. 4b). Similar alterations between genotypes were also observed in females (Fig. S3a-b). Differential expressed gene (DEG) analysis identified 185 significantly changed genes (p ⩽ 0.05) by comparing 5xFAD to Loxp, among which 113 were up-regulated in 5xFAD (including multiple microglial markers with highest fold changes, such as *Trem2, Tyrobp, Cd68, and Clec7a*), and 72 genes were down-regulated (neuronal markers such as *Homer1* and *Fos* were found among the top altered list) (Fig. 4c). A separate DEG comparison of 5xFAD-mKO vs. 5xFAD showed a total of 123 genes significantly altered (p ⩽ 0.05), among which 81 genes were down-regulated (including microglial genes *Trem2, C1qb, Cd68, and Clec7a*), and 42 genes were up-regulated (including *Homer1*) (Fig. 4d). Interestingly, the DEG list generated from 5xFAD vs. Loxp comparison was highly overlapped with the DEG list generated by comparing 5xFAD-mKO to 5xFAD (Fig. S3c). Moreover, among the 113 genes that were induced by Aβ pathology (5xFAD vs. Loxp), 72 were down-regulated by cGAS deletion (5xFAD-mKO vs. 5xFAD) (Fig. 4e). Further query of these 72 genes in WiKiPathway database indicated strong association with microglial pathogen phagocytosis function (Fig. 4f), suggesting that the loss of cGAS greatly diminishes the microglial response to Aβ pathology on a transcription level.

**Figure 4.**
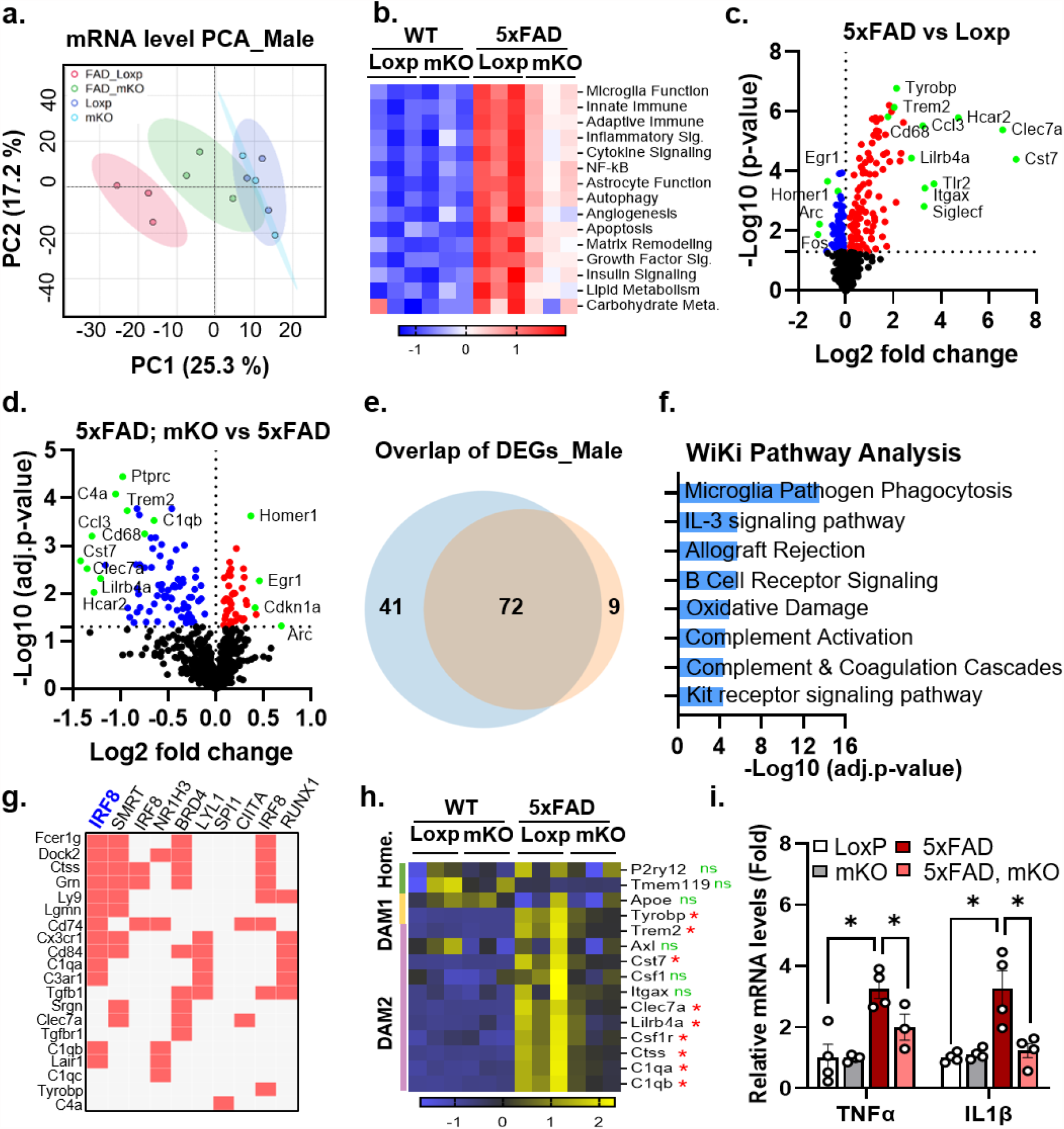
Loss of microglial cGAS ameliorates transcriptomic alterations in 5xFAD mice. a) PCA analysis based on mRNA levels from the NanoString neuroinflammation panel. (n=3/group, male, 7 months old). b) Heatmap displaying pathway scores calculated using NanoString data. c) Volcano plot showing DEGs by comparing 5xFAD to Loxp group. Red and blue color represents increased and decreased expression, respectively. Green color indicates genes of special interest. d) Volcano plot showing DEGs by comparing 5xFAD-mKO to 5xFAD group. e) Venn diagram showing overlap DEGs between plaque-induced genes (5xFAD vs. Loxp, blue color) and genes decreased upon cGAS deletion (5xFAD; mKO vs. 5xFAD, orange color) measured using Nanostring (p-value ⩽ 0.05). f) WiKi Pathway analysis using 72 overlapped gene from e. g) Transcription factor analysis based on ChEA database using 25 overlapped gene from Fig. S3h. h) Microglial gene signature comparison among different genotypes based on NanoString counts. ns or * denotes the significance level by comparing 5xFAD-mKO to 5xFAD group. i) qPCR detection of TNFα and IL1β expression in mouse brain samples (n=4/group, male, 7 months old). ANOVA followed with Tukey’s test for f). *p ⩽ 0.05 and ns (p > 0.05).

Consistently, NanoString analysis in female CRM tissue identified a total of 174 genes that were induced by Aβ pathology (5xFAD compared to Loxp), 56 of which were down-regulated by cGAS knockout (5xFAD-mKO vs. 5xFAD) (Fig. S3d-f). These 56 genes were highly related to microglial phagocytosis and complement function according to WiKiPathway analysis (Fig. S3g). Further examination of the 25 genes that were stimulated by Aβ and suppressed by cGAS deletion in both sexes (Fig. S3h) by a ChEA Transcription Factor Targets analysis revealed that the majority of these genes were downstream targets of the IRF8 transcription factor (Fig. 4g). As a microglia-specific transcription factor in the CNS, IRF8 is critical in regulating microglial activation (30). These findings suggest that the cGAS deficiency-induced decrease in microglial response to the plaque observed in 5xFAD-mKO (compared to 5xFAD) may be mediated by IRF8 down-regulation.

Increased DAM has been well-established as a prominent microglial feature in AD (31). Strikingly, we found that cGAS loss in microglia from 5xFAD background markedly abolished the Aβ pathology-induced DAM markers, while the homeostatic microglial population appears to be intact (Fig. 4h). This suggests that cGAS is a critical modulator for DAM generation under Aβ pathology condition. Consistent with the fact that microglia as a main source of brain cytokines under disease conditions, qPCR analysis of inflammatory gene expression (*TNFα, IL1β*) showed a down-regulation of neuroinflammation upon cGAS deletion (Fig. 4i). Together, these results elucidated that cGAS is indispensable for full activation of DAM phenotype and neuroinflammation in Aβ pathology.

### 2.6 Selective microglial cGAS ablation has neuroprotective effects in Aβ pathology

Mice lacking microglial cGAS exhibited improved spatial memory learning (Fig. 1e-g), suggesting a preserved neuronal functional network under Aβ pathology. These observations prompted us to further explore the mechanism(s) of neuroprotective effect in 5xFAD-mKO mice. Formation of dystrophic neurites is an important feature often seen around plaques (32), which is characterized by enhanced lysosomal marker Lamp1 (33). Interestingly, we detected markedly decreased Lamp1 signal in the hippocampal area of 5xFAD-mKO mice compared to that of 5xFAD (Fig. 5a-5c). Similarly, the number of Lamp1^+^ area and the ratio of the area to the respective total field of view were significantly declined in the cortex of 5xFAD-mKO mice (Fig. 5d-5f). Further, the amount of individual plaque associated-Lamp1^+^ signal was greatly decreased in mice with microglial cGAS deficiency (Fig. 5g and 5h). In parallel with the limited dystrophic neurites, the expression of Homer1, an important scaffolding protein of the post synaptic density (PSD), was largely preserved in 5xFAD-mKO mice brains vs. 5xFAD (Fig. 5i). The neuroprotective effect of cGAS ablation was also evidenced by the elevated PSD95 and Synaptophysin protein levels from brain lysates of 5xFAD-mKO mice (Fig. 5j).

**Figure 5.**
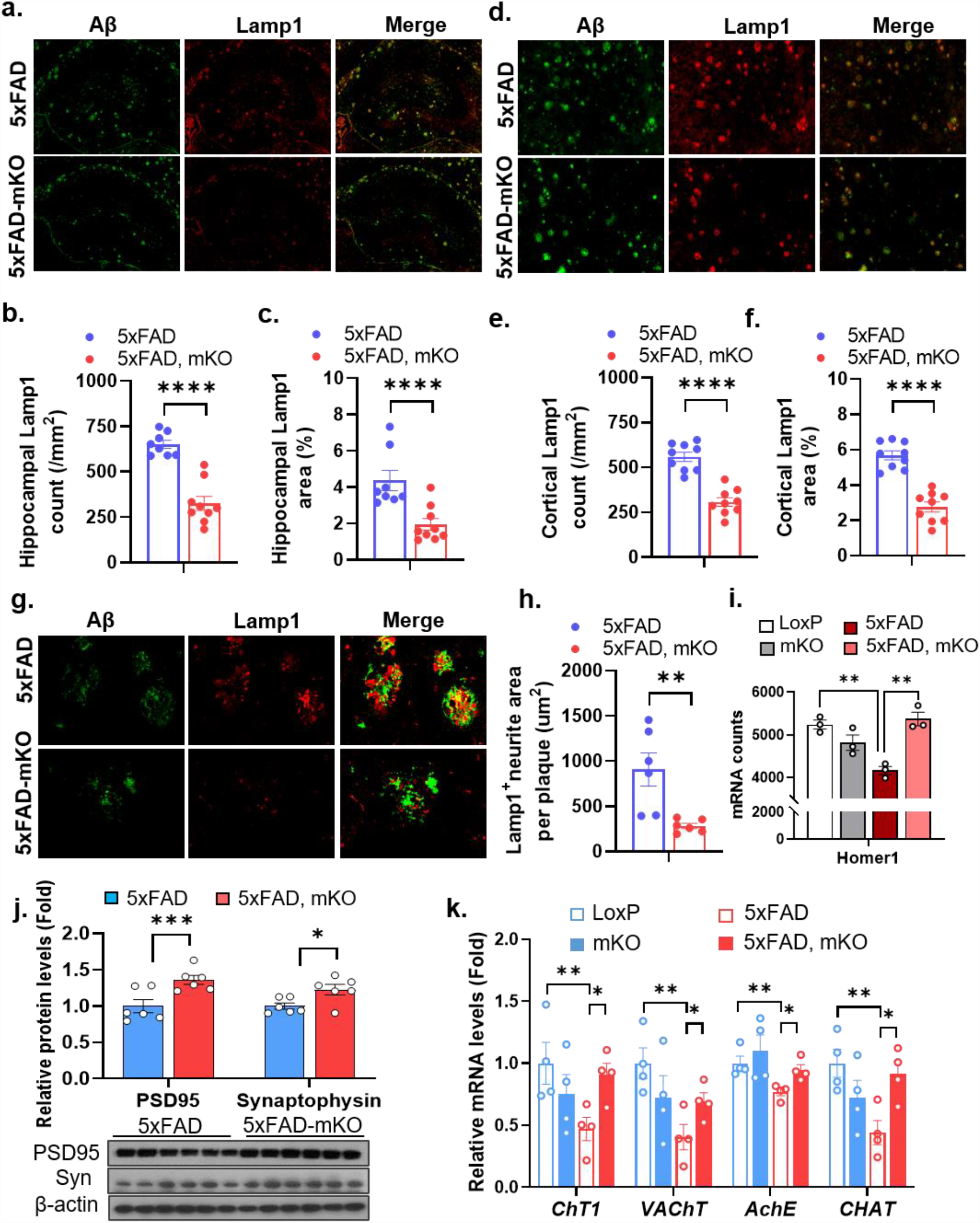
Selective microglial cGAS ablation prevents dystrophic neurites formation. a) Hippocampal IF co-staining using Aβ and Lamp1 antibodies in 7-months-old mice. b) Quantification of overall hippocampal Lamp1 density, and c) percent area ratio in hippocampus region, n=3 mice/group with 3 sections for each mouse. d) Cortical IF co-staining using Aβ and Lamp1 antibody in 7-months-old mice. e) Quantification of overall cortical Lamp1 density, and f) percent area ratio in cortex region, n=3 mice/group with 3 sections for each mouse. g) IF staining, and h) Quantification of plaque-associated Lamp1 levels. i) mRNA counts of Homer1 gene measured with NanoString. n=3/genotype. j) Western-blot, and quantification of PSD95 and synaptophysin protein levels from mouse brain lysates, n=6/genotype. k) qPCR analysis of cholinergic neuron markers expression in brain lysates, n=6/genotype. Student’s t-test for b), c), e), f), h), and j), ANOVA followed with Tukey’s test for i) and k). *p ⩽ 0.05, **p ⩽ 0.01, ****p ⩽ 0.0001, and ns (p > 0.05).

Severe loss of cholinergic neurons was widely observed in AD and has been proposed to contribute to memory and attention deficits (34). Consistently, we detected significant loss of cholinergic neuronal markers in the CRM homogenate of 5xFAD mice compared to wild-type littermates, including choline acetyltransferase (ChAT), vesicular acetylcholine transporter (VAChT), acetylcholinesterase (AChE), and choline transporter (CHT1) (Fig. 5k). Interestingly, deletion of microglial cGAS almost completely rescued the decline of these cholinergic gene expression (Fig. 5k), suggesting preservation of cholinergic production and recycling system in the absence of cGAS. Together, these findings indicate that elevated microglial cGAS may promote cognitive decline through facilitating dystrophic neurite formation and induction of synapse loss as well as disruption of cholinergic neurons in Aβ pathology.

## 3. Discussion

This study emphasized the pivotal roles of microglial cGAS in the development of AD. Conditional deletion of cGAS in microglia limited Aβ deposition, significantly alleviated neuroinflammation, and reduced neuronal damage, which led to better cognitive behavioral outcomes in mice. Previous studies have explored the involvement of cGAS-STING pathway in Aβ and Tau pathology using a mouse model of cGAS whole-body knockout (19, 20). However, given that cGAS is expressed in multiple cell types within the brain (21-24), it would be beneficial and significant to investigate the specific role(s) of cGAS in specific brain cell types. Moreover, components of cGAS-STING pathway are highly expressed in peripheral organs (35), particularly in macrophages (8). Since macrophage infiltration has been implicated in AD pathogenesis (36, 37), it is essential to exclude any potential influence from peripheral cells in understanding the cell-specific role of cGAS in AD. Furthermore, considering that microglia play critical roles in brain development during specific stages (38, 39), avoiding genetic manipulation during these periods may lead to more accurate information when studying an age-related disease such as AD. To address these challenges, the current study employs an inducible, microglia-specific cGAS knockout model, allowing for a better understanding of the role of microglia in AD pathogenesis, and offers new options for the therapeutic targeting of cGAS-STING pathway in AD.

The Aβ peptide deposition-induced formation of extracellular senile plaques is a hallmark of AD (40). Accumulating evidence from genetic, biochemical, and animal model studies has strongly suggested that excessive Aβ accumulation plays a central role in AD pathogenesis, supporting the Amyloid Hypothesis (41-43). In this study, we demonstrated that the absence of cGAS in microglia leads to a reduction in Aβ plaque load, suggesting that cGAS is involved in regulating the balance between Aβ production and clearance. Multiple lines of evidence suggested that microglia play complicated roles in Aβ pathology: Microglia are the major cell type which mediate Aβ clearance through phagocytosis activities (44); conversely, microglia can facilitate propagation of Aβ pathology via seeding (45, 46); studies have also reported that microglia produce a substantial amount of APP, which can be cleaved internally to produce Aβ (47, 48). Modulation of microglia-enriched AD risk genes, such as knockout of Trem2, has shown inconsistent outcomes on amyloid or tau pathologies (49-52). Meanwhile, elevation of Trem2 has different effects at different stages of AD progression. For instance, induction of Trem2 suppresses amyloid accumulation and neuritic dystrophy at the pre-plaque formation stage in 5xFAD mouse model (0-2 months), but it has no impact during plaque accumulating middle to late stages (2-6 months) (53). Collectively, this suggests that the plaque lowering effect of cGAS deletion at the plaque accumulating stage (2-6 months) in our study may be Trem2 independent. Indeed, we identified IRF8 as a potential transcription regulator driving cGAS activation-related microglial reactivity, which was reminiscent of the microglial signature property characterized in human AD (54), further highlighting the potential contribution of cGAS elevation in AD pathogenesis.

In addition to tau neurofibrillary tangles and Aβ plaque, neuroinflammation has been well-recognized as a prominent feature of AD (55). Our previous study showed that loss of sulfatides, a class of myelin-enriched sphingolipid, in the CNS led to an AD-like neuroinflammatory transcriptome, primarily resembling a DAM signature (56). One intriguing direction for future research would be to investigate whether cGAS participates in microglial activation induced by sulfatide loss.

Notably, our NanoString results revealed a significant difference in microglia-specific gene expression between wild-type and cGAS-deficient mouse brains under Aβ pathology. Microglia lacking cGAS exhibited a shifted gene expression profile that forms an intermediate state between homeostatic- and DAM-like states. This is characterized by the preserved elevation of *Apoe, Axl, Csf1*, and *Itgax*, along with the abolished induction of DAM signatures including *Trem2, Clec7a, Csf1r*, and *Lilrb4a*. Moderately activated microglia have been recognized for their neuroprotective effects in the early stages of AD by phagocytosis of Aβ deposits. However, this beneficial function declines upon excessive Aβ accumulation (57, 58). In late stages of AD, overactivated microglia contributes to elevated proinflammatory cytokine production and has reduced Aβ clearance capacity (59). Given the decreased plaque load and inflammatory cytokine levels observed in 5xFAD-mKO mice, it is reasonable to hypothesize that the loss of cGAS leads to a state of moderate microglial activation, which prevents the detrimental consequences of microglial overactivation. Further investigation focusing on a more detailed microglia-specific, temporal transcriptomic analysis and functional study is warranted to test this hypothesis.

We observed a robust protective effect on synaptic integrity in the brain of 5xFAD-mKO mice compared to 5xFAD, as demonstrated by the preserved levels of Homer1, PSD95, and synaptophysin upon cGAS deletion. This is in line with the observation of decreased plaque-associated dystrophic neurites, which reflects the extent of damage to neuronal processes under Aβ-induced neurotoxicity. These alterations may help to maintain neuronal function, thus may be responsible for the improved performance of 5xFAD-mKO mice in cognitive tests. Microglia are known to interact with synapses under both physiological and pathological conditions (60). Removal of excessive synapses by microglia contributes to neuronal maturation during development (61), while microglial synaptic pruning has also been demonstrated to mediate synapse loss in AD (62). Importantly, the complement system plays a critical part in orchestrating this process through the labeling of synapses with C1q and its downstream molecule C3, followed with phagocytic elimination by microglia expressing complement 3 receptor (C3R) (63). Interestingly, multiple complement pathway components, including *C1q, C4a*, and *C3ar1*, were found decreased in 5xFAD-mKO brain vs. 5xFAD, strongly supporting the notion that elevated microglial cGAS drives synaptic loss through up-regulation of complement-mediated synaptic pruning in AD. Thus, further investigation into the glia-neuron interaction under cGAS-deficient condition may elucidate the detailed mechanisms behind the neuroprotective effects of targeting the cGAS-STING pathway in AD.

In summary, our study has shed new light on the functional and mechanistic roles of cGAS in microglia-mediated Aβ accumulation, neuroinflammation, and Aβ pathology-induced synaptic damage. Given the prevalence of AD and its vast impact on human health, our work offers a unique and innovative approach to interrogate the molecular basis of AD pathogenesis. This study could thus hold promise in identifying potential new drug-targeting strategies and could pave the way for new avenues for AD treatment.

## 4. Materials and Methods

### 4.1 Human samples and study approval

Postmortem human brain and spinal cord samples were provided by the NIH NeuroBioBank and Biggs Institute Brain Bank of UTHealth SA. Written consents were obtained for the use of human samples for research.

### 4.2 Animal model

All animal studies were performed in accordance with the guideline approved by the Institutional Animal Care and Use Committee (IACUC) of University of Texas Health San Antonio (UTHSA). Mice were provided with free access to food and water housed under 12/12 h light/dark cycles. Teklad laboratory diet (ENVIGO, Cat. #7012) was used to maintain mouse lines. cGAS fl/fl mouse was generated by flanking exons of the *Mb21d1* gene with LoxP sequences. To generate the microglia-specific *Mb21d1* knockout mouse, cGAS fl/fl mice were crossed with Cx3cr1-CreERT2 mice (Stock No: 020940, the Jackson Laboratory, Bar Harbor, ME, USA). The resulted cGAS fl/fl; Cx3cr1+/- mice were further crossed with 5xFAD (Stock No: 006554, the Jackson Laboratory) to generate 4 genotypes that were used in the current study: Loxp (cGAS fl/fl; Cx3cr1-/-), mKO (cGAS fl/fl; Cx3cr1+/-), 5xFAD (5xFAD; cGAS fl/fl; Cx3cr1-/-), and 5xFAD mKO (5xFAD; cGAS fl/fl; Cx3cr1+/-). All groups were given intraperitoneal injection of tamoxifen (80 mg/kg bodyweight) for 4 consecutive days at 2 months old to induce knockout of *Mb21d1* gene post developmental stage. All animal experiment groups were randomly assigned with mice of desired genotype.

### 4.3 Nest building test

The nest building evaluation was performed according to procedure described in previous report (64) with a 5-point rating scale system. Mice were transferred to individual housing cage at 1 h before dark phase with free access of food and water. An intact piece of cotton sponge of 3 g in weight was provided to each mouse. Rating was performed on the next morning based on the following criteria: 1) the amount and weight of intact sponge (pieces over 0.1 g); 2) the identification of nest site; and 3) nest wall circumference. When the criteria do not agree, a “split of difference” was granted by adding 0.5 in between scores.

### 4.4 Morris water maze test

The maze was set up using a water tank of 120 cm in diameter, in which a circular target platform of 11 cm in diameter was placed within the north-west quadrant. Water was pre-filled to 1 cm above the target platform surface one day before experiment with white-colored paint added to the water. Visual cues were placed around the tank at a height visible from the water level. Mice were acclimated in the behavior room for 1 h daily before trials. For training (spatial acquisition) days, four start positions (including N, S, E, W) were semi-randomized every day, Mice were placed in the water to explore for 1 min for each trial, and was guided to the platform for a 15-second-stay if it did not find the platform within 1 min. Four trials were performed each day per mouse. Probe test was performed 24 h post the last training day, the target platform was removed, each mouse was placed from the south position to complete a 1-min trial in order to assess spatial reference memory.

### 4.5 mRNA extraction and gene expression analysis

Snap frozen cerebrum samples were cryofractured in liquid nitrogen. Approximately 10 mg of tissue per sample was used for extracting mRNA with TRIzol Reagent according to manufacturer’s instruction (Thermo Fisher Scientific, Cat. #15596018). RNA concentration was determined using Spectrophotometer (DeNovix). RNA Integrity (RIN) was determined using the 4150 TapeStation system (Agilent) via RNA ScreenTapes. Subsequent gene expression using NanoString nCounter Technology was performed with Neuroinflammation Panels on nCounter SPRINT Profiler per manufacturer’s instructions (NanoString Technologies). Initial data analysis was performed using nSolver Analysis Software 4.0, which was composed of background thresholding at mean value of Negative Controls, Positive Control Normalization, and CodeSet Content Normalization. Additional analyses including Fold Change analysis between groups and pathway analysis were performed using nSolver Advanced Analysis Software 2.0. For qPCR measurement, 1 μg mRNA per sample was used to perform reverse transcription (Qiagen, Cat. # 205311). Gene expression levels were then detected using SYBR Green (Applied Biosystems, Cat. #A25742) method using primer sequence listed in Supplementary Table 1. Target gene levels were normalized to endogenous housekeeping gene *Gapdh* levels using ΔΔCT method. Data were presented as fold change over respective controls.

### 4.6 Immunoblotting analysis

Frozen samples were cryofractured in liquid nitrogen, 30 mg of tissue per sample was homogenized in N-PER Neuronal Protein Extraction Reagent (Thermo Fisher Scientific, Cat. # 87792) with protease and phosphatase inhibitors (Thermo Fisher Scientific, Cat. # 78442). Samples were incubated on ice for 10 min and centrifuged at 12,000 x g for 15 min at 4 °C. Supernatant was collected for protein quantification using BCA method. 30 μg protein/sample was resolved on NuPAGE 4–12% Bis-Tris gels on the NuPAGE electrophoresis system (Life technologies). Proteins were then transferred onto PVDF membranes, blocked in 1% bovine serum albumin, and incubated with primary antibodies at 4 °C overnight. After washed with TBST, the blots were incubated with horseradish peroxidase-conjugated secondary antibody and developed using chemiluminescence (ECL) method (Thermo Fisher Scientific, Cat. # 32106). A list of antibodies used were included in Supplementary Table 2.

### 4.7 Immunofluorescence staining

Brain tissues were collected upon harvest of animal post saline perfusion, incubated overnight in paraformaldehyde (4% v/v), dehydrated in 10, 20, and 30% sucrose buffer before imbedded into frozen blocks. Serial cross sections (10 μm) of the frozen tissue were collected followed with re-hydration and citric acid-based antigen retrieval. Sections were rehydrated and blocked with 10% goat serum (Sigma, Cat. # G9023) in PBST (PBS containing 0.05% Triton X-100) at room temperature, incubated with primary antibody with 5% goat serum-PBST at 4 °C overnight in a humidified chamber. After washed three times in PBST, slides were incubated with fluorescence-labeled secondary antibody for 1 h at room temperature, before mounted with ProLong™ Diamond Antifade Mountant with DAPI (Thermo Fisher Scientific, Cat. # P36971). Images were collected using a fluorescence microscope (KEYENCE, Cat. # BZ-X800).

### 4.8 Amyloid beta level evaluation

Snap-frozen whole cerebrum tissue was cryofractured in liquid nitrogen. Tissue (25∼30 mg) was used for protein extraction according to protocol described previously (65). Briefly, tissue was homogenized in 15 volumes of Tris-buffered saline (TBS) and centrifuged at 100,000 × g for 1 h at 4 °C using a TLA-55 rotor in an Optima TLX Ultracentrifuge (Beckman Coulter). The first supernatant, TBS-soluble fraction, was frozen in liquid nitrogen and stored at –80 °C. Pellets were washed with 200 μL TBS buffer before re-suspended in 15 volumes (w/v of tissue) of TBSX (TBS buffer containing 1% Triton X-100) and mix gently by rotation at 4 °C for 30 min. After centrifuge at 100,000 × g for 1 h at 4 °C, supernatant was collected as TBSX-soluble fraction. Pellets were re-suspended into 400 μL of 5 M GuHCl, mixed by rotation at room temperature for 6 h, and span at 16,000 × g for 30 min, supernatant was collected as GuHCl-soluble fraction. All buffers used were supplied with protease and phosphatase inhibitors (Calbiochem). Levels of Aβ peptides (amyloid-β1–38, amyloid-β1–40, amyloid-β1–42) in each fraction were measured using V-PLEX Aβ Peptide Panel 1 kit (Meso Scale Discovery, Cat. # K15200E-1) according to manufacturer’s instruction.

### 4.9 Statistics

All data were shown as mean ± standard error of mean (SEM) unless specified. For animal experiments, age- and sex-matched mice were assigned to different treatment groups randomly to prevent potential bias. Data represent results from both male and female unless specified in figure legends. All biochemistry results were representative of at least 3 repeated experiments or as indicated. PCA analysis was performed using MetaboAnalyst 5.0. Statistical analysis was performed using GraphPad Prism 9. Unpaired two-tailed t-test was used for the comparison between two groups. One-way ANOVA followed with Tukey’s test was performed to compare multiple groups. p ⩽ 0.05 was considered statistically significant.

## 5. Declarations

## Acknowledgments

We thank the Healthspan and Functional Assessment Core of UT Health San Antonio for providing equipment and helping with behavior evaluations. We thank the NIH NeuroBioBank and Biggs Institute Brain Bank of UTHealth SA for providing human postmortem frozen tissues for this study.

## Consent for publication

All authors have reviewed the manuscript and reached consent for publication.

## Authors’ contributions

S.H, F.L., and X.H. were involved in the conceptualization and design of the project. S.H., X.L., H.W., N.M., and A.B. performed experiments, and collected and analyzed the data. F.L. contributed to the generation of the mouse model and discussion of the project. S.H. wrote the first draft of the manuscript. S.H., S.Z., F.L., and X.H. contributed to data interpretation, data analysis and editing of the text. X.H. directed the project and provided laboratory resources for the study.

## Data availability statement

Related data that support the findings of this study are available from the corresponding author upon reasonable request.

## References

1. Ho JY, and Franco Y. The rising burden of Alzheimer’s disease mortality in rural America. SSM Popul Health. 2022;17:101052.

2. Griciuc A, and Tanzi RE. The role of innate immune genes in Alzheimer’s disease. Curr Opin Neurol. 2021;34(2):228–36.

3. Karch CM, and Goate AM. Alzheimer’s disease risk genes and mechanisms of disease pathogenesis. Biol Psychiatry. 2015;77(1):43–51.

4. Pimenova AA, Raj T, and Goate AM. Untangling Genetic Risk for Alzheimer’s Disease. Biol Psychiatry. 2018;83(4):300–10.

5. Chen X, and Holtzman DM. Emerging roles of innate and adaptive immunity in Alzheimer’s disease. Immunity. 2022;55(12):2236–54.

6. Sun L, Wu J, Du F, Chen X, and Chen ZJ. Cyclic GMP-AMP synthase is a cytosolic DNA sensor that activates the type I interferon pathway. Science. 2013;339(6121):786–91.

7. Bai J, and Liu F. Nuclear cGAS: sequestration and beyond. Protein Cell. 2022;13(2):90–101.

8. Chen Q, Sun L, and Chen ZJ. Regulation and function of the cGAS-STING pathway of cytosolic DNA sensing. Nat Immunol. 2016;17(10):1142–9.

9. Yang H, Wang H, Ren J, Chen Q, and Chen ZJ. cGAS is essential for cellular senescence. Proc Natl Acad Sci U S A. 2017;114(23):E4612–E20.

10. Gui X, Yang H, Li T, Tan X, Shi P, Li M, et al. Autophagy induction via STING trafficking is a primordial function of the cGAS pathway. Nature. 2019;567(7747):262–6.

11. Li T, and Chen ZJ. The cGAS-cGAMP-STING pathway connects DNA damage to inflammation, senescence, and cancer. J Exp Med. 2018;215(5):1287–99.

12. Wang H, Hu S, Chen X, Shi H, Chen C, Sun L, et al. cGAS is essential for the antitumor effect of immune checkpoint blockade. Proc Natl Acad Sci U S A. 2017;114(7):1637–42.

13. Masanneck L, Eichler S, Vogelsang A, Korsen M, Wiendl H, Budde T, et al. The STING-IFN-beta-Dependent Axis Is Markedly Low in Patients with Relapsing-Remitting Multiple Sclerosis. Int J Mol Sci. 2020;21(23).

14. Standaert DG, and Childers GM. Alpha-synuclein-mediated DNA damage, STING activation, and neuroinflammation in Parkinson’s disease. Proc Natl Acad Sci U S A. 2022;119(17):e2204058119.

15. Yu CH, Davidson S, Harapas CR, Hilton JB, Mlodzianoski MJ, Laohamonthonkul P, et al. TDP-43 Triggers Mitochondrial DNA Release via mPTP to Activate cGAS/STING in ALS. Cell. 2020;183(3):636–49 e18.

16. Barrett JP, Knoblach SM, Bhattacharya S, Gordish-Dressman H, Stoica BA, and Loane DJ. Traumatic Brain Injury Induces cGAS Activation and Type I Interferon Signaling in Aged Mice. Front Immunol. 2021;12:710608.

17. Li Q, Cao Y, Dang C, Han B, Han R, Ma H, et al. Inhibition of double-strand DNA-sensing cGAS ameliorates brain injury after ischemic stroke. EMBO Mol Med. 2020;12(4):e11002.

18. Muhammet F. Gulen NS, Alexander Keller, Marius Schwabenland, Chong Liu, Selene Glück, Vivek V. Thacker, Lucie Favre, Bastien Mangeat, Lona J. Kroese, Paul Krimpenfort, Marco Prinz & Andrea Ablasser cGAS–STING drives ageing-related inflammation and neurodegeneration. Nature. 2023.

19. Xie X, Ma G, Li X, Zhao J, Zhao Z, and Zeng J. Activation of innate immune cGAS-STING pathway contributes to Alzheimer’s pathogenesis in 5xFAD mice. Nat Aging. 2023;3(2):202–12.

20. Udeochu JC, Amin S, Huang Y, Fan L, Torres ERS, Carling GK, et al. Tau activation of microglial cGAS-IFN reduces MEF2C-mediated cognitive resilience. Nat Neurosci. 2023.

21. Wang X, Yang C, Wang X, Miao J, Chen W, Zhou Y, et al. Driving axon regeneration by orchestrating neuronal and non-neuronal innate immune responses via the IFNgamma-cGAS-STING axis. Neuron. 2023;111(2):236–55 e7.

22. Sharma M, Rajendrarao S, Shahani N, Ramirez-Jarquin UN, and Subramaniam S. Cyclic GMP-AMP synthase promotes the inflammatory and autophagy responses in Huntington disease. Proc Natl Acad Sci U S A. 2020;117(27):15989–99.

23. Jeffries AM, and Marriott I. Human microglia and astrocytes express cGAS-STING viral sensing components. Neurosci Lett. 2017;658:53–6.

24. Yu H, Liao K, Hu Y, Lv D, Luo M, Liu Q, et al. Role of the cGAS-STING Pathway in Aging-related Endothelial Dysfunction. Aging Dis. 2022;13(6):1901–18.

25. Zhang Y, Chen K, Sloan SA, Bennett ML, Scholze AR, O’Keeffe S, et al. An RNA-sequencing transcriptome and splicing database of glia, neurons, and vascular cells of the cerebral cortex. J Neurosci. 2014;34(36):11929–47.

26. Zhang Y, Sloan SA, Clarke LE, Caneda C, Plaza CA, Blumenthal PD, et al. Purification and Characterization of Progenitor and Mature Human Astrocytes Reveals Transcriptional and Functional Differences with Mouse. Neuron. 2016;89(1):37–53.

27. Russ DE, Cross RBP, Li L, Koch SC, Matson KJE, Yadav A, et al. A harmonized atlas of mouse spinal cord cell types and their spatial organization. Nat Commun. 2021;12(1):5722.

28. Spangenberg E, Severson PL, Hohsfield LA, Crapser J, Zhang J, Burton EA, et al. Sustained microglial depletion with CSF1R inhibitor impairs parenchymal plaque development in an Alzheimer’s disease model. Nat Commun. 2019;10(1):3758.

29. Thakur S, Dhapola R, Sarma P, Medhi B, and Reddy DH. Neuroinflammation in Alzheimer’s Disease: Current Progress in Molecular Signaling and Therapeutics. Inflammation. 2023;46(1):1–17.

30. Masuda T, Tsuda M, Yoshinaga R, Tozaki-Saitoh H, Ozato K, Tamura T, et al. IRF8 is a critical transcription factor for transforming microglia into a reactive phenotype. Cell Rep. 2012;1(4):334–40.

31. Deczkowska A, Keren-Shaul H, Weiner A, Colonna M, Schwartz M, and Amit I. Disease-Associated Microglia: A Universal Immune Sensor of Neurodegeneration. Cell. 2018;173(5):1073–81.

32. Sadleir KR, Kandalepas PC, Buggia-Prevot V, Nicholson DA, Thinakaran G, and Vassar R. Presynaptic dystrophic neurites surrounding amyloid plaques are sites of microtubule disruption, BACE1 elevation, and increased Abeta generation in Alzheimer’s disease. Acta Neuropathol. 2016;132(2):235–56.

33. Condello C, Yuan P, Schain A, and Grutzendler J. Microglia constitute a barrier that prevents neurotoxic protofibrillar Abeta42 hotspots around plaques. Nat Commun. 2015;6:6176.

34. Ferreira-Vieira TH, Guimaraes IM, Silva FR, and Ribeiro FM. Alzheimer’s disease: Targeting the Cholinergic System. Curr Neuropharmacol. 2016;14(1):101–15.

35. Skopelja-Gardner S, An J, and Elkon KB. Role of the cGAS-STING pathway in systemic and organ-specific diseases. Nat Rev Nephrol. 2022;18(9):558–72.

36. Munoz-Castro C, Mejias-Ortega M, Sanchez-Mejias E, Navarro V, Trujillo-Estrada L, Jimenez S, et al. Monocyte-derived cells invade brain parenchyma and amyloid plaques in human Alzheimer’s disease hippocampus. Acta Neuropathol Commun. 2023;11(1):31.

37. Gate D, Rezai-Zadeh K, Jodry D, Rentsendorj A, and Town T. Macrophages in Alzheimer’s disease: the blood-borne identity. J Neural Transm (Vienna). 2010;117(8):961–70.

38. Paolicelli RC, Bolasco G, Pagani F, Maggi L, Scianni M, Panzanelli P, et al. Synaptic pruning by microglia is necessary for normal brain development. Science. 2011;333(6048):1456–8.

39. Michell-Robinson MA, Touil H, Healy LM, Owen DR, Durafourt BA, Bar-Or A, et al. Roles of microglia in brain development, tissue maintenance and repair. Brain. 2015;138(Pt 5):1138–59.

40. Serrano-Pozo A, Frosch MP, Masliah E, and Hyman BT. Neuropathological alterations in Alzheimer disease. Cold Spring Harb Perspect Med. 2011;1(1):a006189.

41. Hardy J, and Allsop D. Amyloid deposition as the central event in the aetiology of Alzheimer’s disease. Trends Pharmacol Sci. 1991;12(10):383–8.

42. Karran E, and De Strooper B. The amyloid hypothesis in Alzheimer disease: new insights from new therapeutics. Nat Rev Drug Discov. 2022;21(4):306–18.

43. Selkoe DJ. The molecular pathology of Alzheimer’s disease. Neuron. 1991;6(4):487–98.

44. Lee CY, and Landreth GE. The role of microglia in amyloid clearance from the AD brain. J Neural Transm (Vienna). 2010;117(8):949–60.

45. d’Errico P, Ziegler-Waldkirch S, Aires V, Hoffmann P, Mezo C, Erny D, et al. Microglia contribute to the propagation of Abeta into unaffected brain tissue. Nat Neurosci. 2022;25(1):20–5.

46. Venegas C, Kumar S, Franklin BS, Dierkes T, Brinkschulte R, Tejera D, et al. Microglia-derived ASC specks cross-seed amyloid-beta in Alzheimer’s disease. Nature. 2017;552(7685):355–61.

47. Haass C, Hung AY, and Selkoe DJ. Processing of beta-amyloid precursor protein in microglia and astrocytes favors an internal localization over constitutive secretion. J Neurosci. 1991;11(12):3783–93.

48. Chai Q, Jovasevic V, Malikov V, Sabo Y, Morham S, Walsh D, et al. HIV-1 counteracts an innate restriction by amyloid precursor protein resulting in neurodegeneration. Nat Commun. 2017;8(1):1522.

49. Jay TR, Miller CM, Cheng PJ, Graham LC, Bemiller S, Broihier ML, et al. TREM2 deficiency eliminates TREM2+ inflammatory macrophages and ameliorates pathology in Alzheimer’s disease mouse models. J Exp Med. 2015;212(3):287–95.

50. Parhizkar S, Arzberger T, Brendel M, Kleinberger G, Deussing M, Focke C, et al. Loss of TREM2 function increases amyloid seeding but reduces plaque-associated ApoE. Nat Neurosci. 2019;22(2):191–204.

51. Yuan P, Condello C, Keene CD, Wang Y, Bird TD, Paul SM, et al. TREM2 Haplodeficiency in Mice and Humans Impairs the Microglia Barrier Function Leading to Decreased Amyloid Compaction and Severe Axonal Dystrophy. Neuron. 2016;92(1):252–64.

52. Leyns CEG, Ulrich JD, Finn MB, Stewart FR, Koscal LJ, Remolina Serrano J, et al. TREM2 deficiency attenuates neuroinflammation and protects against neurodegeneration in a mouse model of tauopathy. Proc Natl Acad Sci U S A. 2017;114(43):11524–9.

53. Zhao N, Qiao W, Li F, Ren Y, Zheng J, Martens YA, et al. Elevating microglia TREM2 reduces amyloid seeding and suppresses disease-associated microglia. J Exp Med. 2022;219(12).

54. Zhou Y, Song WM, Andhey PS, Swain A, Levy T, Miller KR, et al. Human and mouse single-nucleus transcriptomics reveal TREM2-dependent and TREM2-independent cellular responses in Alzheimer’s disease. Nat Med. 2020;26(1):131–42.

55. Leng F, and Edison P. Neuroinflammation and microglial activation in Alzheimer disease: where do we go from here? Nat Rev Neurol. 2021;17(3):157–72.

56. Qiu S, Palavicini JP, Wang J, Gonzalez NS, He S, Dustin E, et al. Adult-onset CNS myelin sulfatide deficiency is sufficient to cause Alzheimer’s disease-like neuroinflammation and cognitive impairment. Mol Neurodegener. 2021;16(1):64.

57. Sochocka M, Diniz BS, and Leszek J. Inflammatory Response in the CNS: Friend or Foe? Mol Neurobiol. 2017;54(10):8071–89.

58. Zhang G, Wang Z, Hu H, Zhao M, and Sun L. Microglia in Alzheimer’s Disease: A Target for Therapeutic Intervention. Front Cell Neurosci. 2021;15:749587.

59. Hickman SE, Allison EK, and El Khoury J. Microglial dysfunction and defective beta-amyloid clearance pathways in aging Alzheimer’s disease mice. J Neurosci. 2008;28(33):8354–60.

60. Wolf SA, Boddeke HW, and Kettenmann H. Microglia in Physiology and Disease. Annu Rev Physiol. 2017;79:619–43.

61. Schafer DP, Lehrman EK, and Stevens B. The “quad-partite” synapse: microglia-synapse interactions in the developing and mature CNS. Glia. 2013;61(1):24–36.

62. Hong S, Beja-Glasser VF, Nfonoyim BM, Frouin A, Li S, Ramakrishnan S, et al. Complement and microglia mediate early synapse loss in Alzheimer mouse models. Science. 2016;352(6286):712–6.

63. Wilton DK, Dissing-Olesen L, and Stevens B. Neuron-Glia Signaling in Synapse Elimination. Annu Rev Neurosci. 2019;42:107–27.

64. Deacon RM. Assessing nest building in mice. Nat Protoc. 2006;1(3):1117–9.

65. Youmans KL, Leung S, Zhang J, Maus E, Baysac K, Bu G, et al. Amyloid-beta42 alters apolipoprotein E solubility in brains of mice with five familial AD mutations. J Neurosci Methods. 2011;196(1):51–9

