## Supplementary Information for "Microglial cGAS deletion protects against amyloid-β induced Alzheimer’s disease pathogenesis"

[1]Barshop Institute for Longevity and Aging Studies, University of Texas Health San Antonio; San Antonio, TX 78229, USA. [2]Division of Endocrinology, Department of Medicine, University of Texas Health San Antonio; San Antonio, TX 78229, USA. [3]Metabolic Syndrome Research Center, The Second Xiangya Hospital of Central South University, Changsha, Hunan 410011, China. [4]Division of Diabetes, Department of Medicine, University of Texas Health San Antonio; San Antonio, TX 78229, USA.

Feng Liu,;

**This file includes:**

Supporting text

Figures S1 to S3

Tables S1 to S2

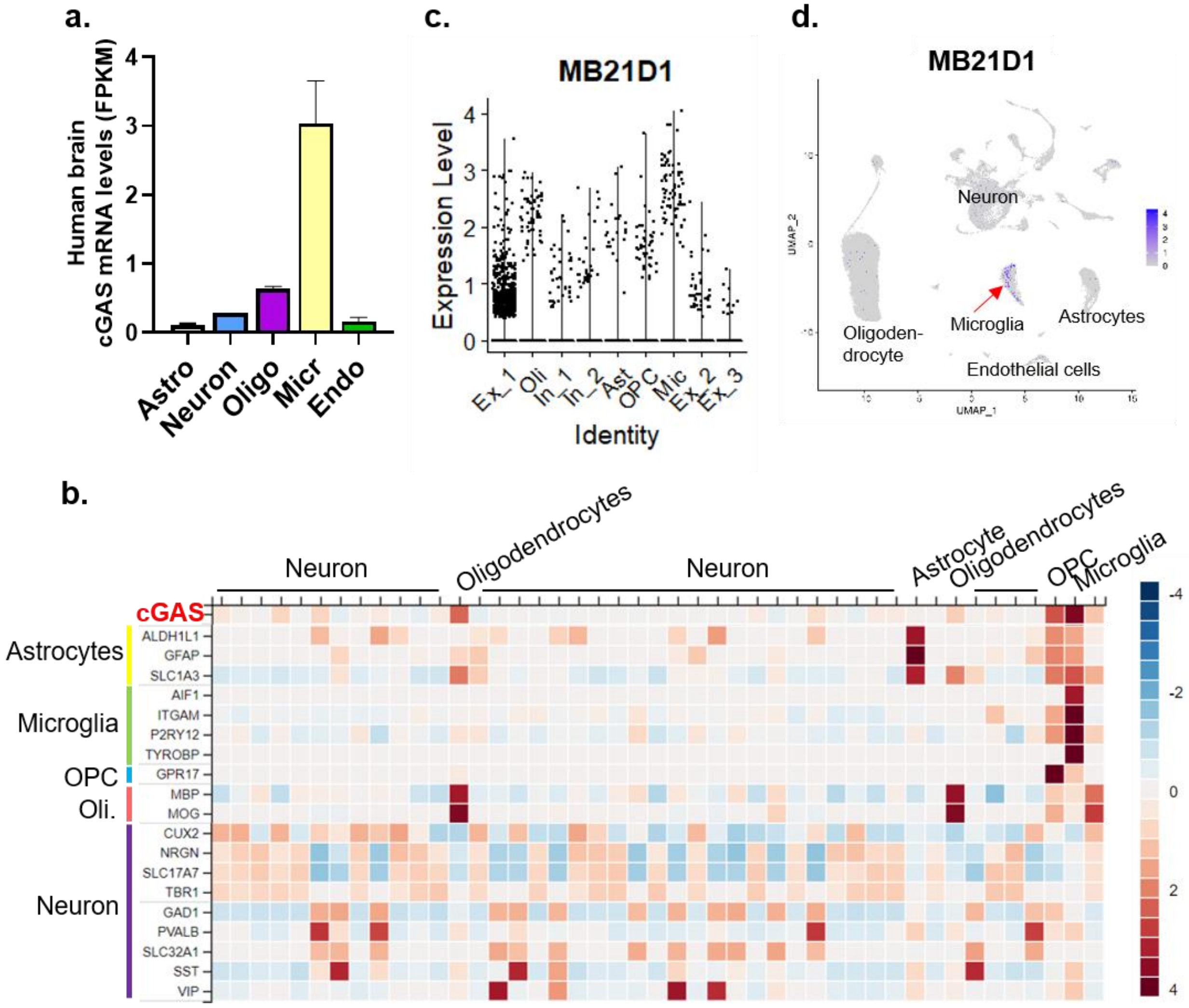


**Figure S1. cGAS expression in human brain and spinal cord tissues.** a) cGAS expression shown by RNA-Seq of cells isolated from human brain, data adapted from (https://www.brainrnaseq.org/). b) cGAS expression in human brain tissue measured by single cell sequencing. Data adapted from “Human Protein Atlas proteinatlas.org”. c) cGAS mRNA levels in different cell populations measured by single-nucleus sequencing, data adapted from (DIO: 10.1038/s41586-019-1195-2). Note that Ex-1 was the most enriched cell population, while microglia had the highest cGAS levels. d) Single cell sequencing database inquiry of cGAS expression in human spinal cord tissues. Data adapted from (https://seqseek.ninds.nih.gov/spinalcordinjury) (DIO: 10.1038/s41467-021-25125-1).

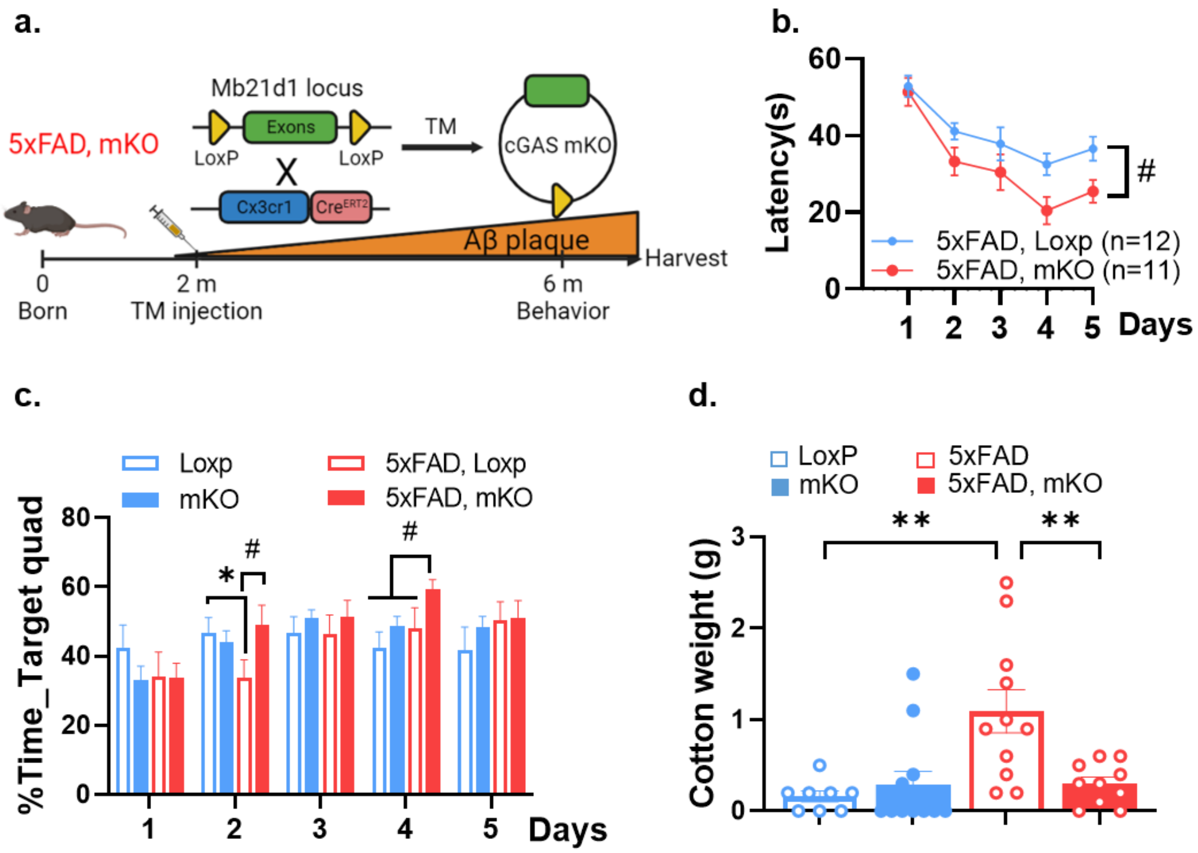


**Figure S2. Behavial phenotypes of microglia-specific cGAS knockout mice.** a) Schematic showing the strategy for establishing the cGAS mKO mouse model. (Figure created using BioRender). b) Latency, and c) Percent time in target quadrants during training in a MWM test. (Showing female data from 6 months old, n=9-12/genotype). d) Weight of un-shredded cotton after a nest building test. (Male and female, 7 months old, n=8-12/group).

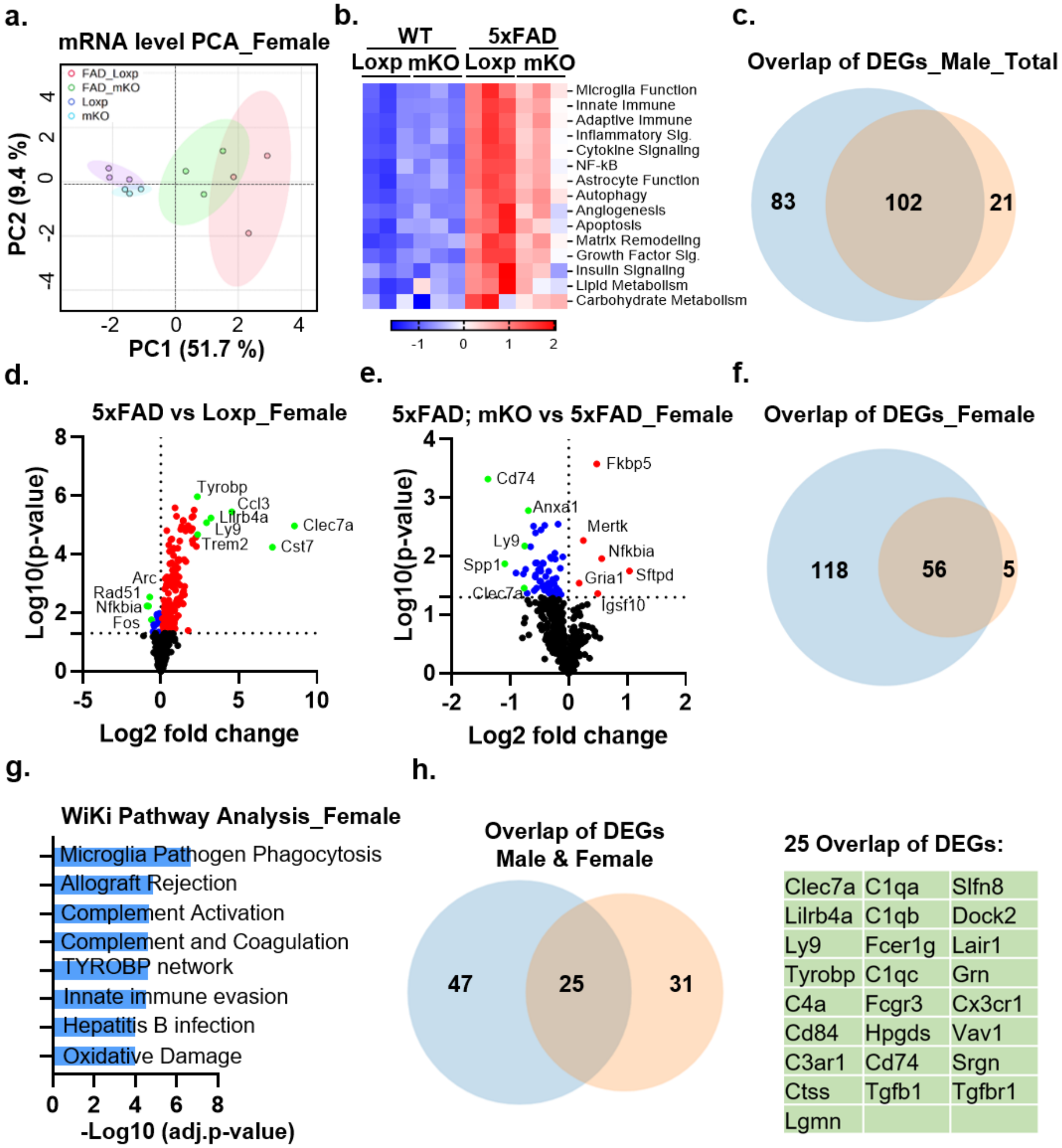


**Figure S3. Transcriptomic alterations in female 5xFAD-mKO mice.** a) PCA analysis based on mRNA levels from NanoString neuroinflammation panel. (n=3/group, female, 7 months old). b) Heatmap displaying pathway scores calculated using female NanoString data. c) Venn diagram showing overlap of DEGs that were altered by Aβ pathology and by cGAS deletion in male mice (note this includes both increased and decreased genes). d) Volcano plot showing female DEGs by comparing 5xFAD to Loxp group. Red and blue color represents increased and decreased expression, respectively. Green color indicates genes of special interest. e) Volcano plot showing female DEGs by comparing 5xFAD-mKO to 5xFAD group. f) Venn diagram showing overlap female DEGs between plaque-induced genes (5xFAD vs. Loxp, blue color) and genes decreased upon cGAS deletion (5xFAD; mKO vs. 5xFAD, orange color) measured using Nanostring (p-value $\leqslant$ 0.05). g) WiKi Pathway analysis using 56 overlapped genes from f. h) Venn diagram and list of the 25 overlapped DEGs that were induced by Aβ pathology and reduced by cGAS deletion in both sexes.

**Table S1. List of primers used**

| Primer name | 5'-sequence-3' |
|---|---|
| **m-GAPDH_F** | GAGAAACCTGCCAAGTATG |
| **m-GAPDH_R** | GGAGTTGCTGTTGAAGTC |
| **m-cGas_F** | GGCAGCTACTATGAACATGTG |
| **m-cGas_R** | CTCAGCGGATTTCCTCGTGGAA |
| **m-IL1b-F** | GAAATGCCACCTTTTGACAGTG |
| **m-IL1b-R** | TGGATGCTCTCATCAGGACAG |
| **m-TNFα_F** | GCCTCTTCTCATTCCTGCTT |
| **m-TNFα_R** | CTCCTCCACTTGGTGGTTTG |
| **m-ChAT_F** | GAGCGAATCGTTGGTATGACAA |
| **m-ChAT_R** | AGGACGATGCCATCAAAAGG |
| **m-ChT1_F** | GCAGCTTTTGGGTGCCTG |
| **m-ChT1_R** | TGTGGAAGCTCCAATAGCTCC |
| **m-VAChT_F** | GGGTCGGCTCGGTCAATC |
| **m-VAChT_R** | CAAATAGCACGCCTATCTTCACAT |
| **h-GAPDH_F** | CATGTTCCAATATGATTCCACC |
| **h-GAPDH_R** | CTCCATGGTGGTGAAGACGC |
| **h-cGas_F** | GGGAGCCCTGCTGTAACACTTCTTAT |
| **h-cGas_R** | CCTTTGCATGCTTGGGTACAAGGT |

**Table S2. List of antibodies used**

| Antibody | Source | Identifier |
|---|---|---|
| **cGAS (mouse)** | CST | 31659S |
| **cGAS (human)** | CST | 15102 |
| **STING** | CST | 13647S |
| **Aβ (MOAB)** | Millipore | MABN254 |
| PU.1 | CST | 2258S |
| **Iba1(immunoflourescence)** | FUJIFILM Wako | 019-19741 |
| **Lamp1** | Thermo Fisher | 14-0112-82 |
| **PSD95** | CST | 3450T |
| **Synaptophysin** | CST | 36406T |
| **β-actin** | CST | 4970S |
